# A novel method to detect intracellular metabolite alterations in MCF-7 cells by doxorubicin induced cell death

**DOI:** 10.1101/812255

**Authors:** Ajay Kumar, Sheetal Patel, Devyani Bhatkar, Nilesh Kumar Sharma

## Abstract

Metabolic reprogramming within cancer cells is suggested as a potential barrier to chemotherapy. Additionally, metabolic tumor heterogeneity is one of factor behind discernible hallmarks such as drug resistance, relapse of tumor and the formation of secondary tumors. In this paper, cell based assays including PI/annexin V staining and immunoblot assay were performed to show the apoptotic cell death in MCF-7 cells treated with DOX. Further, MCF-7 cells were lysed in hypotonic buffer and whole cell lysate was purified by a novel and specifically designed metabolite (100 to 1000 Da) fractionation system as vertical tube gel electrophoresis (VTGE). Further, purified intracellular metabolites were subjected to identification by LC-HRMS technique. The authors show the presence of cleaved PARP 1 in MCF-7 cells treated with DOX. Concomitantly, data show the absence of active caspase 3 in MCF-7 cells. Novel findings are to identify key intracellular metabolites assisted by VTGE system that include lipid (CDP-DG, phytosphingosine, dodecanamide), non-lipid (N-acetyl-D-glucosamine, N1-acetylspermidine and gamma-L-glutamyl-L-cysteine) and tripeptide metabolites in MCF-7 cells treated by DOX. Interestingly, the authors report a first evidence of doxorubicinone, an aglycone form of DOX in MCF-7 cells that is potentially linked to the mechanism of cell death in MCF-7 cells. This paper reports on novel methods and processes that involve VTGE system based purification of hypotonically lysed novel intracellular metabolites of MCF-7 cells treated by DOX. Here, these identified intracellular metabolites corroborate to caspase 3 independent and mitochondria induced apoptotic cell death in MCF-7 cells.

**SIGNIFICANCE STATEMENT:** Metabolic reprogramming in cancer cells is implicated in various tumor hallmarks. Interestingly, thousands of research have addressed the molecular basis of drug treatment and resistance in chemotherapy. But, there is a significant gap in the precise methodologies and approaches in addressing intracellular metabolite alterations. This paper reports on a novel approach that helped reveal new findings on intracellular metabolite changes in case of doxorubicin (DOX) induced cell death in MCF-7 cells. This paper highlights the additional insights on debatable findings available in literature in the contexts of DOX induced cell death mechanisms. In this paper, novel and specifically designed vertical tube gel electrophoresis (VTGE) system is claimed to purify intracellular metabolites and this method is compatible with other biological system.

## INTRODUCTION

According to the Global Cancer Report issued by the World Health Organization, there are over 10 million new cases of cancer each year and over 6 million annual deaths from the disease (1). Breast cancer is reported to affect around 1.6 million women every year throughout the globe and further estimated to increase to 2.2 million cases yearly by 2025.

There are various therapeutic avenues, including chemotherapy, hormonal therapy, radiation therapy and immunotherapy therapy with limited success due to intra- and inter-tumor heterogeneity and adaptive tumor microenvironment within breast tumor (2–8). These therapies are faced with discernible widely accepted problems, including, drug resistance, concomitant its strong side effects such as neurotoxicity, nephrotoxicity, gastrointestinal toxicity vomiting and ototoxicity, recurrence of cancer and poor quality of life post treatment (2–4).

Doxorubicin (DOX), an anthracycline antibiotic is the most commonly anticancer drug to treat broad spectrum of cancer types including breast cancer, bladder cancer, Kaposi’s sarcoma, non-Hodgkin’s and Hodgkin’s lymphoma and acute lymphocytic leukemia (9–12). There are numerous papers that have established the mechanisms of action of DOX including as an inhibitor of topoisomerase II enzyme and via free radical mechanisms that damage, major macromolecules including DNA and proteins that eventually lead to cancer cell death (9–12). In spite of numerous papers that address the molecular mechanisms of DOX an anticancer drug, there is a lack of clarity in the role of DOX to show caspase 3 dependent or independent apoptotic cell death in MCF-7 cells (13–16). Here, there is an emphasis in the literature on the molecular identity of MCF-7 cells that lack active caspase 3 expression and at the same time cleaving of poly (ADP-ribose) polymerase (PARP) 1 due to DOX toxicity (13–16). However, existing facts on DOX induced death of MCF-7 cells are refutable and debatable. There is a lack of evidence on the presence of DOX or aglycone metabolite of DOX (doxorubicinone) in the intracellular compartment and ensuing metabolic changes in MCF-7 cells. Normal cells, such as muscle cells, liver cells are reported to metabolize DOX into doxorubicinone that contributes to cell. Interestingly, the role of doxorubicinone and its presence in MCF-7 cells are not reported till date. Furthermore, there is a limited evidence on the metabolic adaptations in cancer cells including MCF-7 due to DOX mediated toxicity (17–20).

Based on the above problems and gaps in the molecular understanding on DOX mediated MCF-7 cell death, we attempt to show the first evidence on the presence of doxorubcinone, a DOX aglycone metabolite in the intracellular components of MCF-7 cells. Further, our finding provides a clear profile on DOX mediated intracellular metabolite alterations in MCF-7 cells by using a novel VTGE system that precisely purifies metabolites, including organic acids, lipids, amino acids, and tripeptides and further characterized by LC-HRMS techniques.

## MATERIAL AND METHODOLOGY

### Materials

Cell culture reagents were purchased from Invitrogen India Pvt. Ltd. and Himedia India Pvt. Ltd. MCF-7 cells were procured from the National Centre For Cell Science (NCCS), Pune, India. DMSO, Doxorubicin, DMEM High Glucose media, acrylamide and other chemicals were of analytical grade and obtained from Himedia India Pvt. Ltd and Merck India Pvt. Ltd.

### Cell Line Maintenance

The MCF-7 cells were cultured and maintained in DMEM (Dulbecco’s Modified Eagles Medium) (Himedia) with high glucose supplemented with 10% heat inactivated FBS/penicillin (100 units/ml)/streptomycin (100 µg/ml) at 37°C in a humidified 5% CO2 incubator.

### Trypan blue dye exclusion assay

MCF-7 cells were plated onto six well plate (1.5*10^5^ cells per well) and incubated in the presence of complete fresh DMEM high glucose medium supplemented with 10% FBS. After 16-18 h, cells were treated with 2 ml of complete fresh DMEM medium and treated with DOX 100 nM final concentration. A negative control of MCF-7 cells treated with DMSO solvent is designated as DMSO control throughout this paper for intracellular metabolite analysis and comparison. After 48 h of incubation, the media was aspirated to recover the floating dead cells. Next, cells were washed with PBS and treated with 0.25% trypsin-EDTA for 2-3 min at 37°C. Fresh media were added into each well and cells were harvested properly and ensured that earlier collected dead cells are pooled together. Next, the total and viable MCF-7 cell count were performed as per the manufacturer instruction for a routinely used Trypan blue dye exclusion assay (21).

### Propidium iodide/Annexin V staining assay

MCF-7 cells were seeded in duplicates into six well plate at the plating density of 15X10^4^ cell per well. After 16-18 h of plating, MCF-7 cells were with DOX (100 nM) and DMSO as solvent and incubated for 48 h in the presence of a complete culture medium. At the end of incubation, MCF-7 cells were collected by routine trypsinization and cell counting was performed using haemocytometer. Cell suspensions were centrifuged at 1500 rpm for 3 min. Finally, pellets were resuspended in 1 ml of cold PBS buffer. Next, MCF-7 cells were centrifuged and Annexin V binding buffer was added to the pellet. We performed this assay using Annexin V/FITC apoptosis detection kit (ThermoFisher) to stain MCF-7 cells for Annexin V and propidium iodide (PI). Tubes were kept for incubation at room temperature in the dark for 15 min and the annexin binding buffer was added to all the tubes. Then, cells were analysed on BD FACSJazz Cytometer to count the live and dead cells. At least, 10,000 events were collected and analyzed per measurement (21, 22).

### Immunoblotting as an apoptosis assay

To determine the nature of apoptotic cell death pathway, whole cell lysates of MCF-7 cells prepared by RIPA buffer based cell lysis were separated on 10% polyacrylamide gel and then transferred to nitrocellulose membrane. Further, as per the manufacturer instructions from Apoptosis Western Blot Cocktail (pro/p17-caspase-3, cleaved PARP1, muscle actin) (ab136812) kit, we performed immunoblot to detect the presence of active caspase 3 and cleaved PARP 1 in MCF-7 cells treated by DOX and DMSO control (21).

### Preparation of hypotonic whole cell lysate

A question is pertinent to ask the nature of alterations of intracellular metabolites, including lipids, organic acids and tripeptides in MCF-7 cells treated by DOX compared to DMSO control. The detailed methods and processes are given in published Indian Patent Application Number: 201921000760 (21, 23, 24). In brief, DOX and DMSO treated MCF-7 cells for 48 hr cells were placed in hypotonic buffer (10 mM KCl, 10 mM NaCl, 20 mM Tris, pH 7.4) for swelling to facilitate mechanical homogenization. For one million of MCF-7 cells, 500 µl of hypotonic buffer is used. Then, MCF-7 cells were lysed by the help of 31 gauge needle and followed by mechanical homogenization. Further, mechanically lysed MCF-7 cells were centrifuged at 12000 g for 30 min to get clear supernatant of MCF-7 cells treated by DOX and DMSO. Finally, hypotonically lysed whole cell lysate of MCF-7 cells was used for intracellular metabolite purification and identification.

### Purification of intracellular metabolites by VTGE

Next, 250 µl of clear hypotonically lysed whole cell lysates of MCF-7 cells was further diluted to 750 µl with the help of hypotonic buffer. Finally, 750 µl of hypotonically prepared intracellular MCF-7 cell lysate mixed with 250 µl of 4X loading buffer (0.5 M Tris, pH 6.8 and Glycerol) and loaded on a specifically designed VTGE system (Figure 1) that used 15% acrylamide gel (acrylamide:bisacrylamide, 30:1) as matrix to separate molecules based on size. The fractionated elute was collected in electrophoresis running buffer that contains water and glycine and excludes traditional SDS, and other reducing agents. Intracellular metabolite derived from MCF-7 cells were eluted in the same running buffer, which is referred as elution buffer. After loading of intracellular MCF-7 cell lysates with loading buffer, the power supply was connected and voltage and current ratio were maintained to generate 1500-2500 mW of power to achieve the electrophoresis of intracellular lysate including metabolite and large size biomolecules such as proteins. The total run time has been allowed for 2 hr and at the end of 2 hr, lower collecting buffer was collected in tube for further characterization (23, 24). At the end of the run, inner tube containing polyacrylamide gel was removed and processed for coomassie brilliant blue dye staining to ensure that protein components of cell lysate were trapped in the PAGE gel. Further, these eluted metabolite fractions of cell lysates were stored at -20°C for LC-HRMS analysis and identification of metabolites.

**Figure 1.**
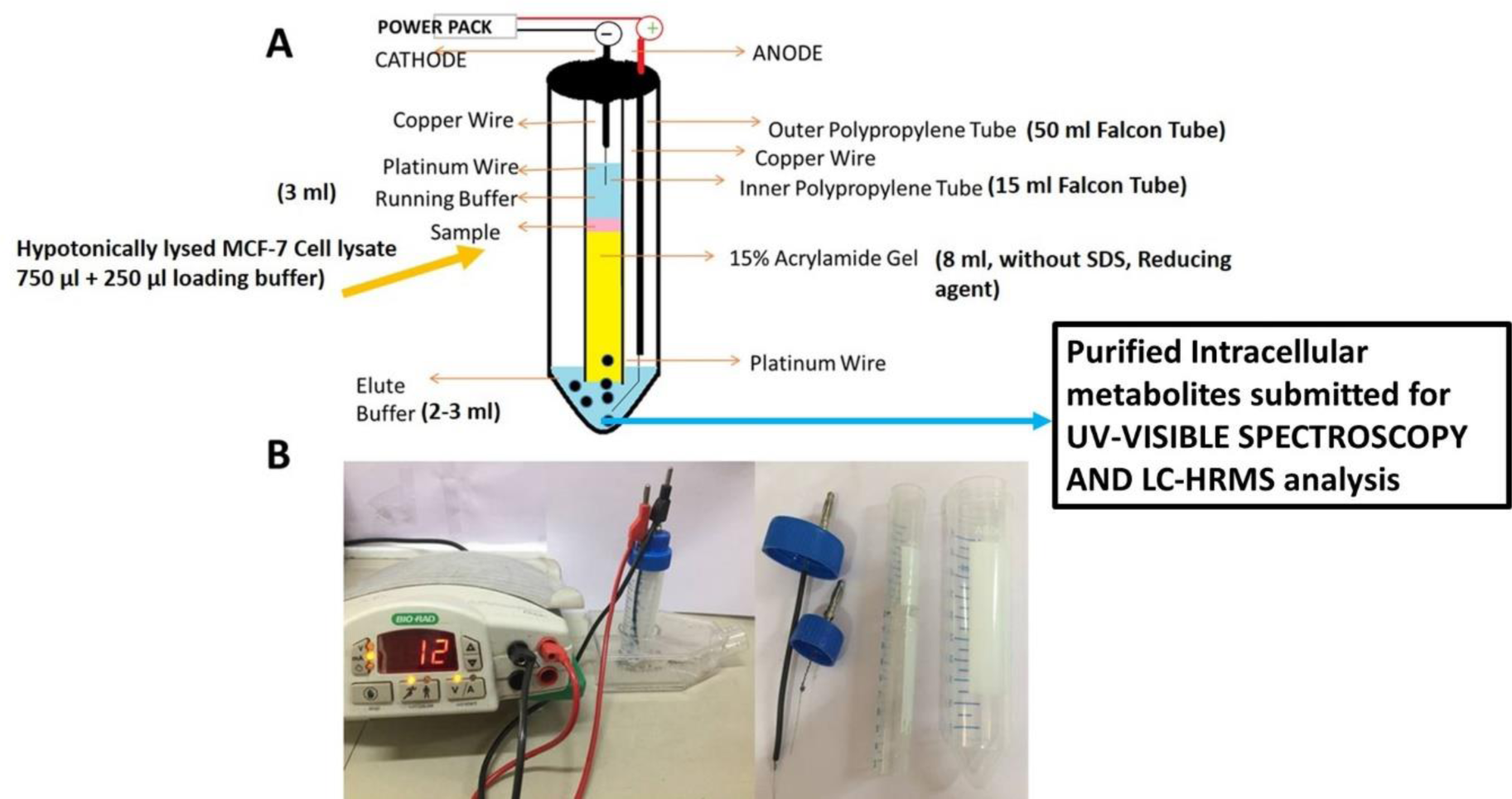
A flow diagram of novel and specifically designed vertical tube gel electrophoresis (VTGE) system for intracellular metabolite purification.

### LC-HRMS intracellular metabolite profiling

The eluted intracellular metabolite of MCF-7 cells in water: glycine buffer from above methods and processes was submitted to LC-HRMS based identification of potential metabolites such as organic acids, lipids, amino acids, and tripeptides. In this method, Agilent TOF/Q-TOF Mass Spectrometer station with an ion source Dual AJS ESI was used for MS/MS analysis. In short, during LC-HRMS, an LC column RPC18 Hypersil GOLD C18 100 x 2.1 mm-3 µm was used and data were acquired in positive mode. During LC, an injection volume of 3.00 µl per sample was used and flow of sample at 0.300 ml/min was maintained. Two solvent channels as 100% Water (0.1% FA in water) and 100% Acetonitrile (90% ACN +10% H2O+ 0.1% FA) was used from 95% and 5% combination. Data was acquired in positive ion mode with a MS/MS absorbance threshold at 5, MS relative threshold at (0.01%) and MS absorbance threshold at 200 (23, 24).

## STATISTICAL ANALYSIS

Data shown are presented as the mean ± SD of at least three independent experiments; differences are considered statistically significant at P < 0.05, using a Student’s t-test.

## RESULTS AND DISCUSSION

The power of metabolic adaptations of cancer cells in response to drug insults including DOX therapeutic is a major hurdle in the success and failure of cancer therapy (17-20, 22, 25, 26). Therefore, a better understanding at intracellular levels of targeted cancer cells in various metabolic pathways, including glucose, amino acid, lipid, and nucleosides will help to understand the cell death pathways and associated drug resistance consequences.

### Dox-induced apoptotic cell death in MCF-7 cells

In the light of molecular basis of DOX effects upon MCF-7 cells, we revisited the cell based assay to demonstrate loss of cell viability and induction of apoptotic cell death in MCF-7 cells at 100 nM dose of DOX. By employing Trypan blue dye exclusion assay, data on total reduction of MCF-7 cells and loss of viability of MCF-7 cells treated with DOX at 100 nm for 48 hr is given Figure 2A, 2B and 2C. Data presented as photomicrographs taken after the treatment of DOX clearly demonstrate the loss of viability of MCF-7 cells up to 95%. However, reduction in total MCF-7 cells are noticed up to 36%. In short, observations suggest that MCF-7 cells are undergoing cellular adaptations to undergo cell death. To address the nature of cell death in MCF-7 cells by DOX, we present data collected from flow cytometry based PI and Annexin V stained MCF-7 cells (Figure 3). As seen earlier, DOX brings highly noticeable loss of viability and similarly, apoptotic assay results show the presence of 50% of MCF-7 cells with apoptosis that is as per the expectations and matches with existing evidence. Further, whole cell lysates of MCF-7 cells treated by DOX are evaluated for the apoptotic markers such as PARP 1 cleavage by using western blot based apoptotic assay kit. An immunoblot photograph shows the clear presence of the cleaved PARP 1 band in case of DOX treated MCF-7 cells over DMSO treated MCF-7 cells (Figure 4). At the same time, there is no detection of active caspase 3 activity by immunoblotting in MCF-7 cells. Taken together, immunoblot data indicate the generation of cleaved PARP 1 due to the DOX toxicity in MCF-7 cells. Lack of active caspase 3 activity is in line with earlier findings. Collectively, different cell based data converge to suggest the presence of apoptotic cell death in MCF-7 cells due to DOX activity and possibly caspase 3 enzyme independent cleavage of PARP 1.

**Figure. 2.**
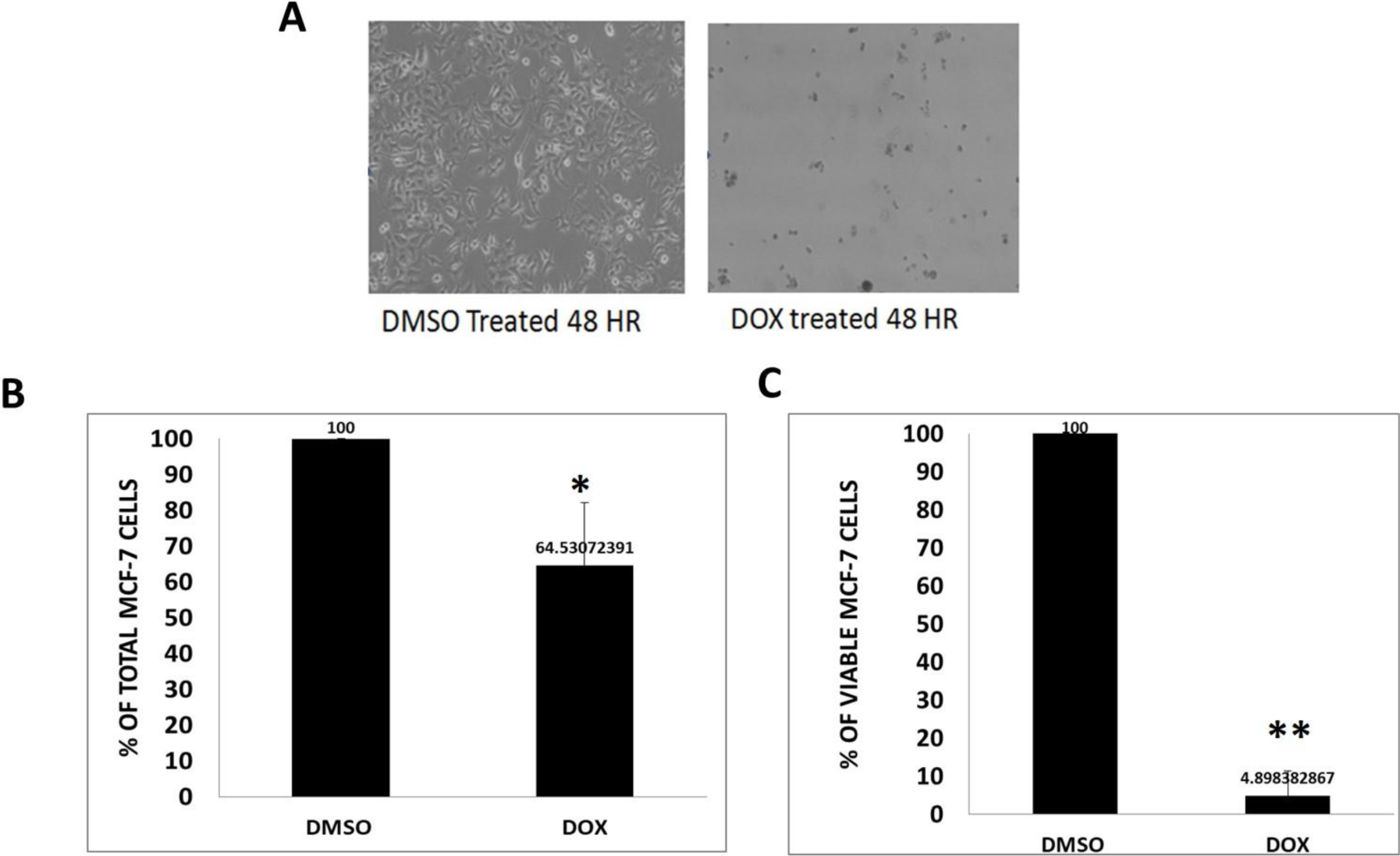
DOX-induced loss of cell viability in MCF-7 cells MCF-7 cells were treated by 100 nm DOX and a solvent control DMSO for 48 hr. Then, harvested cells were subjected to Trypan blue dye exclusion assay to estimate total cell and viable cells. Figure (2A). This figure represents the phase contrast microscopy photographs (100 X magnification) of MCF-7 cells treated with DMSO and DOX for 48 hr. Figure (2B). This bar graph shows total cell count performed by Trypan blue dye exclusion assay on MCF-7 cells treated DMSO and DOX. Figure (2C). This bar graph illustrates the percentage loss of cell viability of MCF-7 cells estimated by Trypan blue dye exclusion assay. Data are represented as mean ± SD. Each experiment was conducted independently three times. The bar graph without an asterisk denotes that there is no any significant difference compared to DMSO control. * Significantly different from DMSO control at the P-value < 0.05. ** Significantly different from DMSO control at P-value < 0.001.

**Figure. 3.**
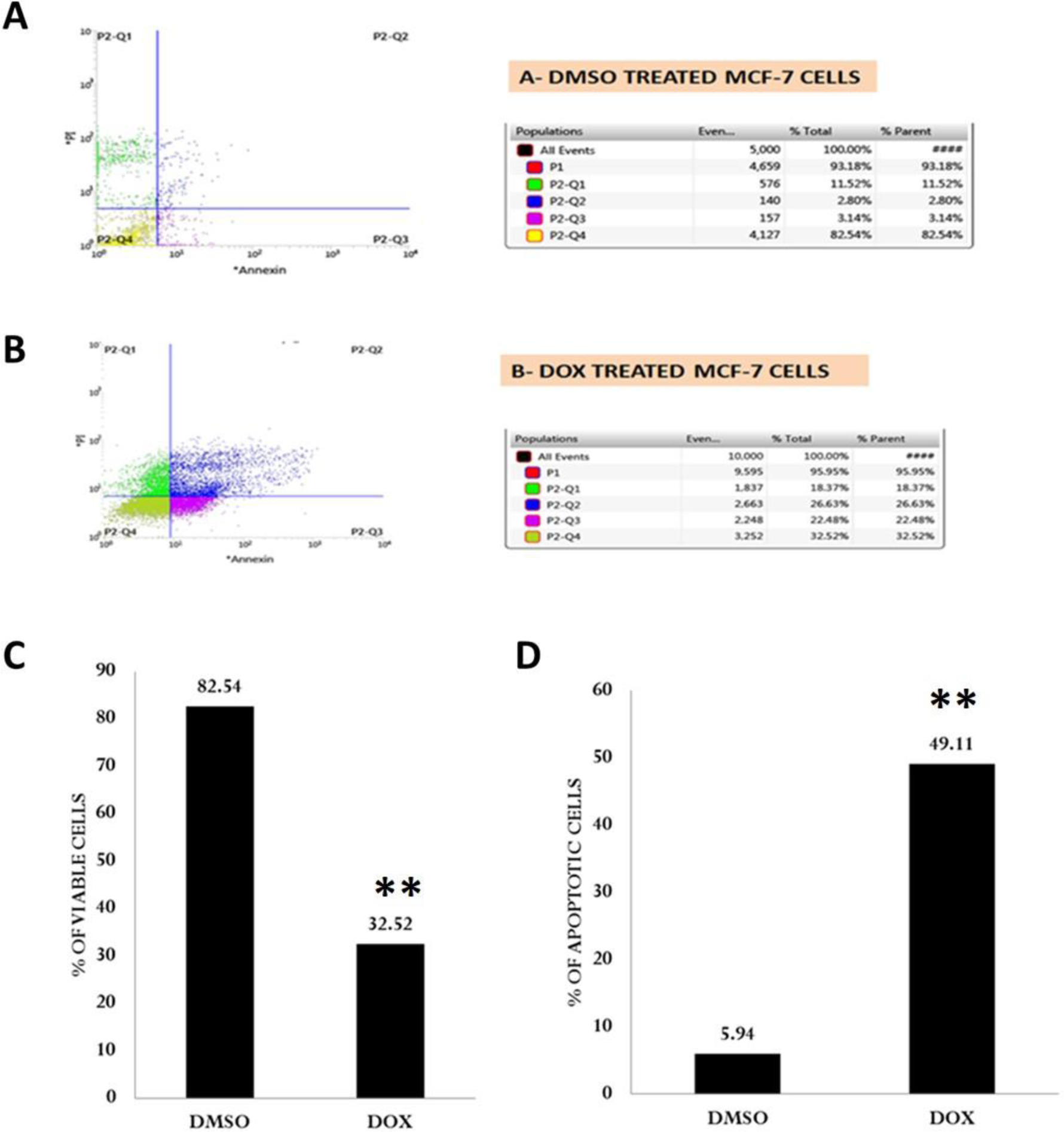
Induction of apoptosis by DOX in MCF-7 cells as determined by PI/annexin V staining In this assay, MCF-7 cells were treated as detailed in Figure 2 legend. At the end of treatment, harvested MCF-7 cells were subjected to PI/annexin V staining and analyzed by flow cytometer. Data represent the cells stained with PI and Annexin V conjugated with FITC for the analysis of apoptotic cells in MCF-7. The scatter plot of cells treated with DMSO and DOX is shown Figure 3A and Figure 3B, respectively. Figure 3C depicts the bar graph representations of loss of cell viability. Figure 3D indicates the percentage of apoptotic MCF-7 cells treated with DMSO and DOX. Data are represented as mean ± SD. Each experiment was conducted independently three times. The bar graph without an asterisk denotes that there is no any significant difference compared to DMSO control. * Significantly different from DMSO control at the P-value < 0.05. ** Significantly different from DMSO control at P-value < 0.001.

**Figure. 4.**
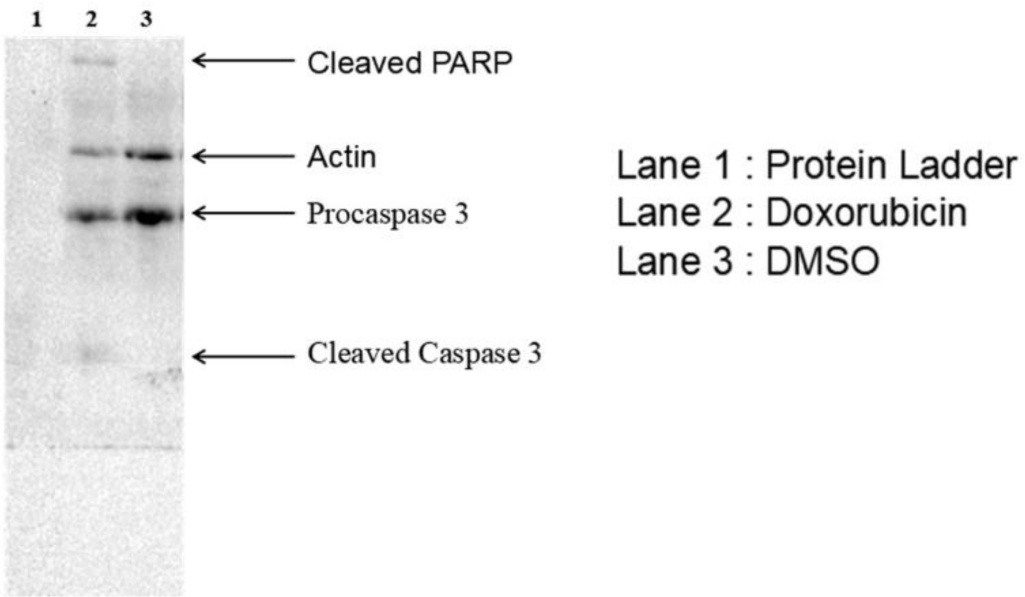
DOX activates cleavage of PARP 1 and absence of activated caspase 3 by immunoblotting Here, MCF-7 cells were treated as detailed in Figure 2 legend. At the end of treatment, harvested MCF-7 cells were submitted for whole cell lysate preparation. Then, 40 µg of proteins were loaded into 10% polyacrylamide gel and were immunoblotting with primary antibody cocktails as per the manufacturer instructions of Apoptosis Western Blot Cocktail (pro/p17-caspase-3, cleaved PARP1, muscle actin) (ab136812) kit. Presented immunoblot were developed by using chemiluminescent substrate and HRP enzyme system.

There is a clear consensus is that DOX is a non-intercalating drug that produces a single- and double-strand break in DNA of treated cancer cells (11, 12, 17-20, 22, 25, 27). Further, the presence of single- and double-strand breaks in DNA is suggested to be mediated by the blockade of topoisomerase II enzyme activity in DOX targeted cancer cells. At preclinical and clinical levels, findings on mode of action by DOX at the molecular level remains debatable. Among several molecular pathways, the role of DOX is reported as the poison of topoisomerase II, generate DNA adduct, formation of reactive oxygen species and intercalative drug that directly binds to DNA (9-12, 17, 22, 25, 27). Furthermore, the role of DOX is suggested to directly impinge upon mitochondria that induce caspase 3 independent death in target cancer cells including MCF-7 cells (13–16). There is well-accepted fact that the evidence on caspase 3 mediated apoptotic cell death in MCF-7 cells is controversial and lack clear understanding.

Among various controversial issues, MCF-7 cells are suggested to lack caspase 3. Interestingly, MCF-7 cells are known to express caspase 7 and further this caspase 7/8 enzyme participates in mitochondria mediated apoptotic cell death (13–16). Evidences suggest that PARP 1 is a substrate for both caspase 3 and caspase 7 (13–16). In this way, the presence of cleaved PARP 1 in MCF-7 treated with DOX may signify the involvement of caspase 7 and mitochondria dependent apoptotic cell death.

A study on effects of metaformin upon MCF-7 cells suggested presence of cleaved PARP1 and involvement of caspase 7 and mitochondria in bringing apoptotic cell death (28). Since MCF-7 cells are deficient of caspase 3 and in line with that we also report on the presence of the active cleaved form of PARP 1. Therefore, a strong suggestion emerges that other form of caspase enzyme as caspase 7/8 may be involved in the DOX mediated MCF-7 cell death. Another way is that deregulation of mitochondria may trigger the formation of cleaved PARP 1. In this direction, the presence of doxorubicinone in the intracellular compartment of MCF-7 cells treated with DOX is one of the interesting observation discussed in this paper. Hence, there is a possibility that mitochondria may be targeted due to the abundance of doxorubcinone and that may trigger PARP 1 mediated apoptotic cell death as evident from PI and Annexin V staining assay. There are clear evidences that support the role of a key nuclear enzyme PARP 1 in apoptosis. Besides main executioner of apoptosis as caspase-3, caspase-7 is shown to cleave PARP1 and Hsp90 by using unique key lysine residues (K(38)KKK) present in the N-terminal proteolytic domain of caspase-7 (16). Further data suggest the role of caspase 7 to cleave PARP1 during VP-16 mediated apoptotic cell death in HL60 and caspase 3 deficient MCF-7 cells (29).

### Total ion chromatogram (TIC) of VTGE system purified intracellular metabolites

Based on our observation and data from earlier findings, caspase 3 independent cleavage of PARP 1 and involvement of mitochondrial metabolic adaptations has not been addressed with reference to DOX toxicity in MCF-7 cells. In view of intracellular metabolic adaptations, there is a need to revisit changes in MCF-7 cells treated by DOX that is lacking in this area of research. The authors highlight the contribution of VTGE system based intracellular metabolite preparation that precisely excludes high molecular weight and complex macromolecules including proteins, complex sugars and nucleic acids and retains only metabolites of M.W. less than 1000 Da. As described in the method section, intracellular metabolite elutes were identified by LC-HRMS technique. A representative TIC is presented for intracellular metabolites of DMSO and DOX treated MCF-7 cells in Figure 5A and 5B, respectively. The authors point out that different types of metabolites, including lipid, amino acids derived, polyamine, simple sugars and tripeptides are detected in the TIC of intracellular metabolite and precisely the M.W of these metabolites ranges from 119 Da. to 841 Da. In turn, data from TIC clearly support and validate the working efficiency of VTGE system claimed in this paper to facilitate identification of intracellular metabolites from in vitro studies similar to present DOX mediated metabolic adaptations in MCF-cells. The present intracellular metabolite profiling is supported by limited attempts in the literature and those approaches have employed different methodology and technique (5, 8, 17, 30, 31).

**Figure. 5.**
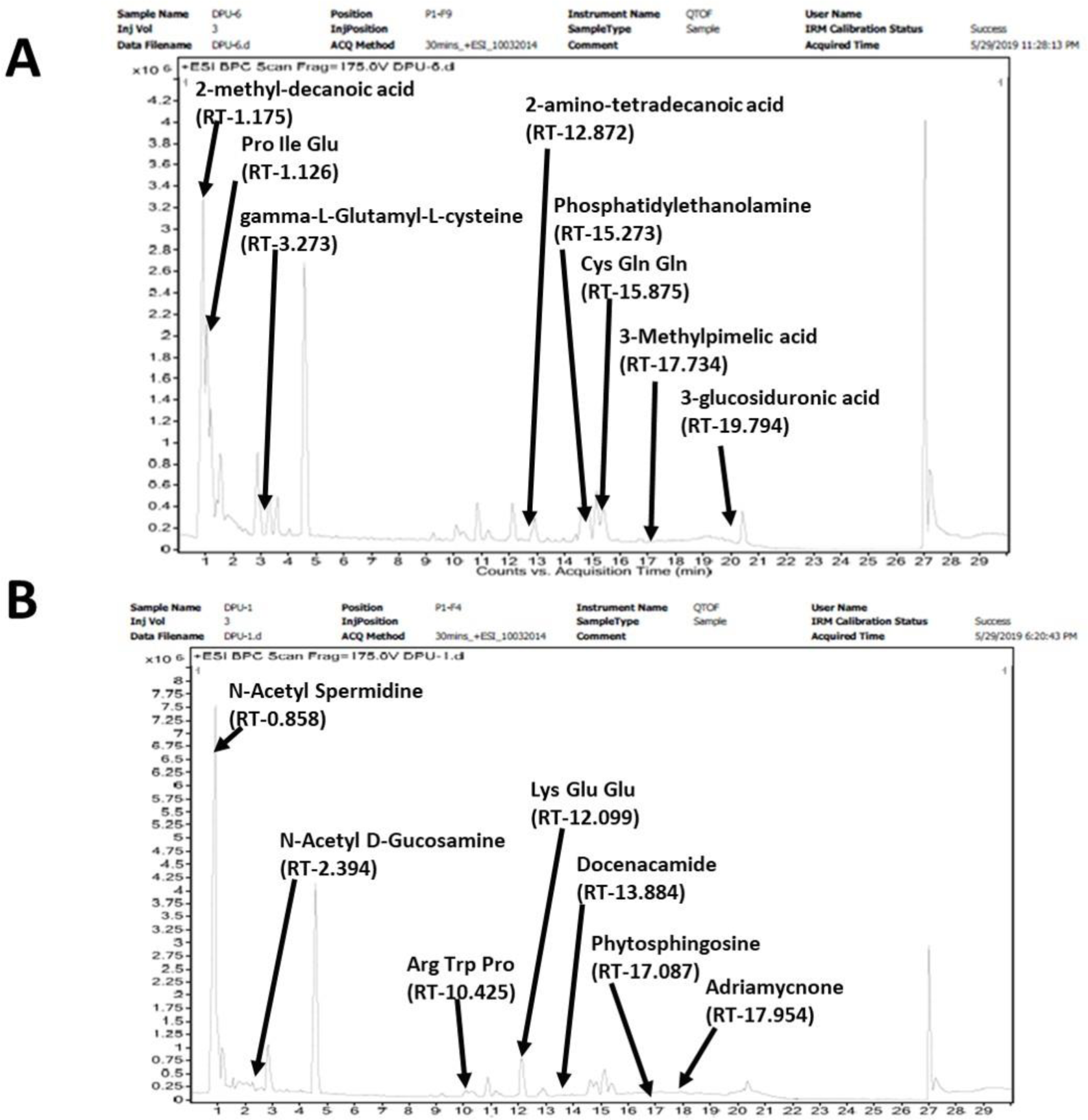
VTGE purified intracellular metabolites identified by LC-HRMS show unique TIC in DOX treated MCF-7 cells To identify intracellular metabolite changes, MCF-7 cells were treated as detailed in Figure 2 legend with DOX and DMSO. Further, MCF-7 cells were hypotonically lysed and whole cell lysate were purified by specifically designed VTGE system. Next, purified intracellular metabolites of MCF-7 cell lysate was identified by LC-HRMS. A total ion chromatogram (TIC) of intracellular metabolite of DOX and DMSO treated MCF-7 cells are given in Figure 5A and 5B, respectively.

### Validation of intracellular doxorubicinone in DOX treated MCF-7 cells

In the past decades, a plethora of research papers reported on DOX and involved molecular pathways that contribute towards sensitivity and resistance in various types of cancer cell lines including MCF-7 cells (17-20, 22, 25-27). But, an interesting observation needs attention that none of published papers have shown a clear evidence on the presence of DOX or metabolized aglycone DOX as doxorubicinone. Data is available to suggest the metabolic conversion of DOX to DOX semiquinone free radicals inside targeted cancer cells, but a clear presence and detection of such DOX or DOX derived free radical by metabolomic approach is lacking. Besides DOX semiquinone free radicals, the conversion of DOX into doxorubicinone by deglycosidation is speculated inside targeted cancer cells. However, not a single paper has addressed the detection of doxorubicinone in MCF-7 cells treated with DOX. Further detection of doxorubicinone in MCF-7 cells will bring additional evidence of the activity of DOX in cells and also may put a step forward in this field that faced a debatable and controversial molecular basis of DOX action to show apoptosis in MCF-7 cells.

Therefore, the authors address the presence of DOX or doxorubicinone in the intracellular component of MCF-7 cells treated DOX by employing a novel VTGE process combined with LC-HRMS techniques for the metabolites profiling. An extracted MS/MS ion spectra is presented in Figure 6 that clearly convince the presence of doxorubicinone in the intracellular compartment of MCF-7 cells treated by DOX. In this paper, we present the first report on the presence of doxorubicinone with a m/z value of 419.081and M.W. 414.103 as DOX derived metabolites that actually got attention in normal cell toxicity and associated side effects in the human body. Metabolism of DOX in biological system including hepatocytes, muscle cells, cancer cells is suggested to generate primary metabolite such as DOX aglycone and adriamycinol by drug metabolizing enzymes (10). Further, the role of DOX derived metabolites such as doxorubicinone is suggested to contribute for dysfunction of mitochondria and release of cytochrome C (32). A recent study suggests the potential binding ability of DOX and aglycone doxorubicinone to the minor groove of DNA (26).

**Figure. 6.**
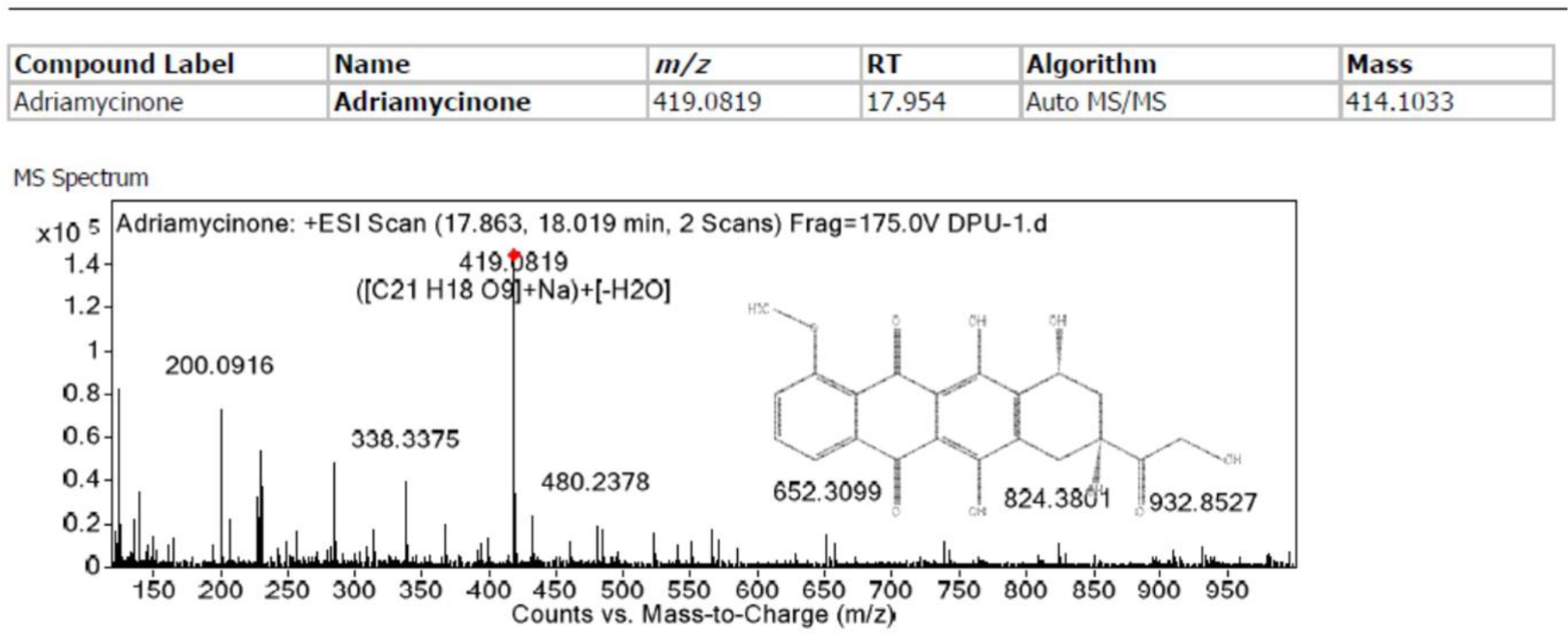
A MS/MS ion spectra of DOX metabolized to doxorubicinone, aglycone form of DOX Among the intracellular metabolite pool, DOX treated MCF-7 cells shows first and novel evidence and identification of doxorubicinone, a glycone product of DOX (Figure 6). In literature, various cell and animal model suggested the presence of metabolized form of DOX. Surprisingly, none of research paper has actually shown the presence of doxorubicinone in MCF-7 cells treated with DOX and as well in other cancer types.

In literature, DOX is known to contain an aglyconic component doxorubicinone and sugar moieties. The aglycone form of DOX as doxorubicinone contains quinone-containing tetracyclic ring with a short side chain containing a carbonyl group at C-13 and a primary alcohol at C-14. Further, daunosamine, a sugar moiety is attached to the tetracyclic ring with the help of glycosidic bond to C-7 position (27). An earlier paper indicated the presence of doxrubicinone, aglycone form of DOX in the plasma of human treated with DOX (11). However, an evidence on the presence of doxorubicinone in the intracellular compartment of MCF-7 is of first report. This report provides additional insight to propose a role of doxorubicinone as a disruptor of mitochondrial function in MCF-7 cells, which are most commonly reported for normal cell toxicity.

Such action of doxorubicinone may be a factor behind toxicity and proapoptotic effects of DOX upon normal cells and suggest mechanisms of action of DOX other than inflicted by the semiquinone free radicals. It is important to highlight that the role of DOX aglycone metabolites are more studied with reference to side effects of DOX in normal cells. However, the potential role of doxorubicinone in cancer cells including MCF-7 cells are not addressed due to controversies in the DOX mediated molecular mechanisms that lead to apoptosis in MCF-7 cells lacking caspase 3 (14, 33). Therefore, this paper proposes to look into another dimension of apoptotic effects of DOX upon MCF-7 cells via the generation of doxorubicinone and subsequent damage to mitochondria and then activation of caspase 7 and finally cleavage of PARP 1.

Further, an argument may be raised that the role of DOX to induce apoptosis in normal cells and MCF-7 cells are different. In this direction, earlier data indicated that DOX has distinct mechanisms to induce apoptosis and this is discriminated due to the p53 activation and free radical exposure (34). The present paper claims that in spite of distinct apoptotic mechanisms of DOX between normal cells to MCF-7 cells, a possible cell death pathway that creates an axis of doxorubicinone mediated mitochondrial dysfunction, release of cytochrome C, cleavage of PARP 1 via caspase 7 and finally apoptotic cell death of MCF-7 that is independent of caspase 3 may be further warranted.

In spite of these existing data, the effects of DOX in MCF-7 cells are delineated at the level of caspase dependent or caspase independent apoptotic cell death, ROS mediated cell death and mitochondrial mediated cell death. However, a simple evidence is missing in this area of research that may be pertinent to ask the intracellular evidence on DOX or doxorubicinone in treated cancer cells including MCF-7 cells. Based on the literature, there is not a single paper that attempts to address by that may provide evidence on the presence of DOX or DOX metabolized product such as aglycone DOX doxorubicinone intracellular content of MCF-7 cells. Based on the literature, it appears that in all in vitro studies involving MCF-7 and DOX, emphasis is placed onto molecular mechanism that mediate apoptotic cell death due to DOX. However, evidence on the presence of DOX or DOX metabolized product as doxorubucinone is not addressed and reported at the intracellular level.

### Lipid profile alterations in DOX treated MCF-7 cells

Based on the literature and our data, the role of DOX to induce apoptosis in MCF-7 is suggested to involve mitochondrial deregulation and alterations in the lipid metabolism (15, 17-20, 35–38). Therefore, we evaluated lipid profiles in the intracellular compartment of MCF-7 cells treated DOX over DMSO control. Additionally, there is a lack on the report that delineates precise details on the changes in the intracellular metabolite components in MCF-7 cells treated by DOX. Because, metabolic adaptations by MCF-7 cells and other cancer cells in response to DOX or other anticancer drugs are key cellular changes that confers sensitivity or resistance to a particular drug. In another way, revelations on the changes in the intracellular metabolite changes in MCF-7 cells treated by DOX may provide additional platform to understand anticancer molecular pathways. On the other hand, same approaches may be employed to observe intracellular metabolite changes in case of DOX resistant MCF-7 cells and target amenable metabolic pathways that create hindrances in achieving better DOX responses.

Identifying intracellular lipid metabolites in MCF-7 cells are assisted by VTGE system and LC-HRMS techniques and details of lipid metabolites are given in Table 1 and Figure 7. In a detailed analysis of lipid metabolites observed in DOX and DMSO treated MCF-7 cells, the data clearly suggest that the precise nature of lipid as phytosphingosine, dodecanamide and cytidine diphosphate diacylglycerol (CDP-DG) are elevated in DOX treated MCF-7 cells that shows cell death over DMSO control. Here, the authors interpret observations towards doxorubicinone mediated deregulation of mitochondrial lipid metabolisms that resulted in distinct lipid metabolite abundance in DOX treated MCF-7 cells over DMSO control.

**Figure. 7.**
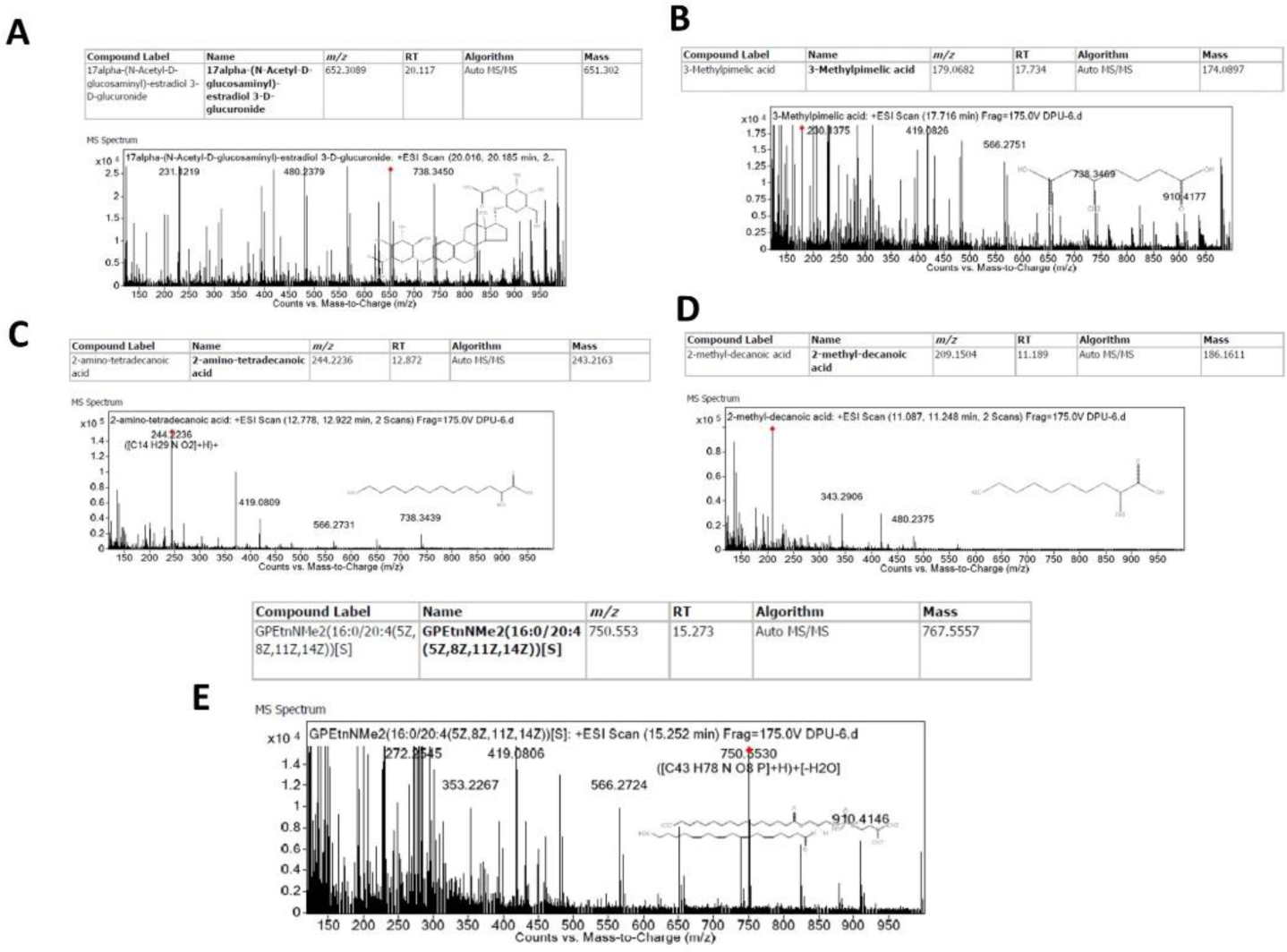
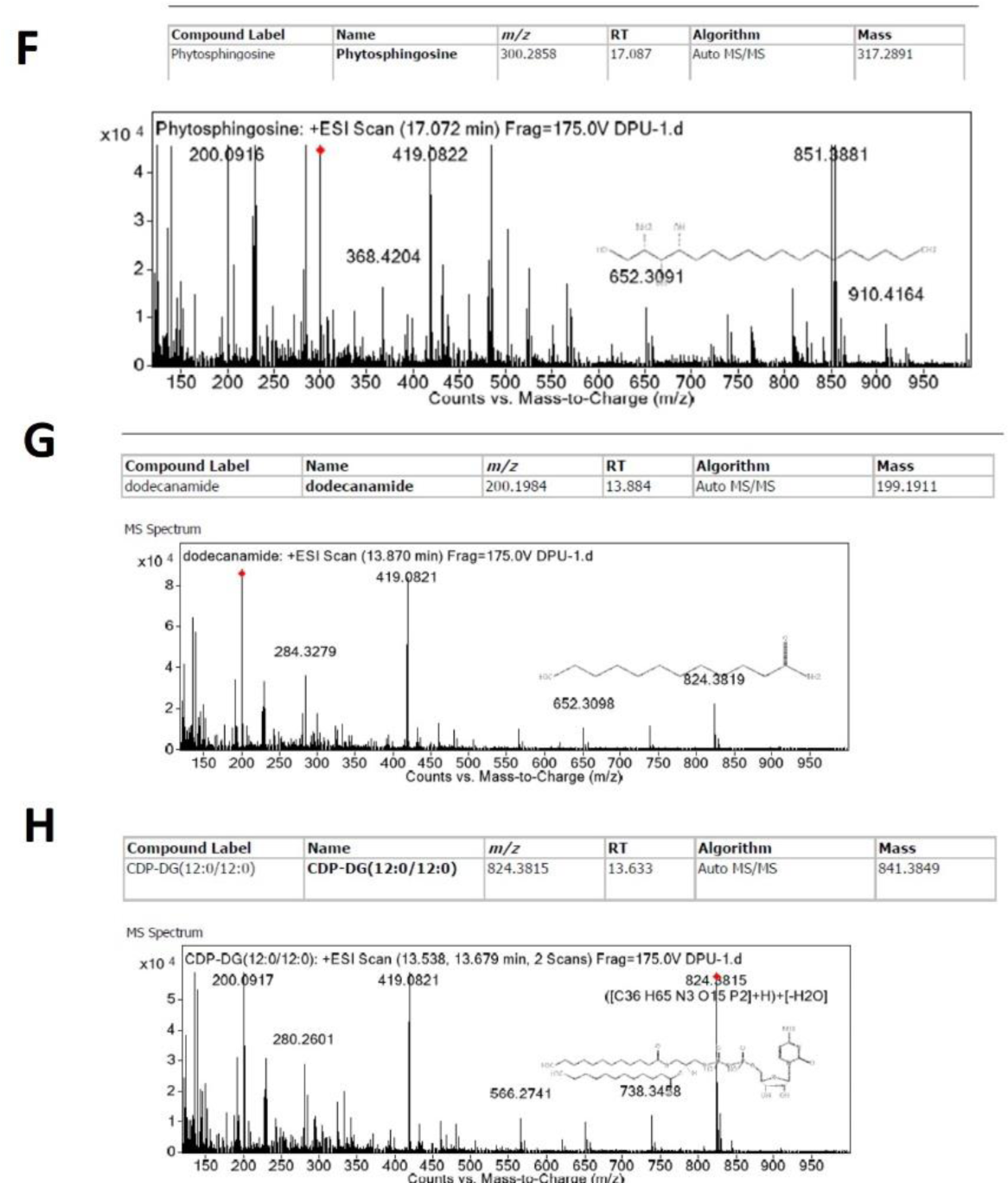
DOX treated MCF-7 cells displays novel intracellular lipid metabolite alterations To identify lipid metabolite alterations due to DOX, MCF-7 cells were treated as detailed in Figure 2 legend with DOX and DMSO. Further, MCF-7 cells were hypotonically lysed and whole cell lysate were purified by specifically designed VTGE system. Based on LC-HRMS identification and analysis, data suggest CDP-DG, Phytosphingosine and dodecanamide as potential lipid metabolite accumulation in DOX treated MCF-7 cells (Figure 7F-H). On the other hand, DMSO control treated MCF-7 depicts a distinct set of lipid profiles, including 17α-(N-acetyl-D-glucosaminyl) estradiol 3-glucosiduronic acid, 2-amino-tetradecanoic acid, 2-methyl-decanoic acid, 3-Methylpimelic acid and Phosphatidylethanolamine over DOX treated MCF-7 cells (Figure 7A-E).

**Table 1:** A list of intracellular lipid metabolite comparative abundance in DOX over DMSO treated MCF-7 cells.

| Sr. No | Name of Lipids | DMSO treated MCF-7 cells lysate | DOX treated MCF-7 cells lysate | RT | m/z | M.W. |
| --- | --- | --- | --- | --- | --- | --- |
| 1 | Phytosphingosine | Not detectable | 45529 | 17.087 | 300.285 | 317.289 |
| 2 | Dodecanamide | Not detectable | 87514 | 13.884 | 200.198 | 199.191 |
| 3 | 17α-(N-acetyl-D-glucosaminyl) estradiol 3-glucosiduronic acid | 26545 | Not detectable | 20.117 | 652.308 | 651.301 |
| 4 | 2-amino-tetradecanoic acid | 155340 | Not detectable | 12.872 | 244.223 | 243.216 |
| 5 | 2-methyl-decanoic acid | 100812 | Not detectable | 1.175 | 209.150 | 186.161 |
| 6 | 3-Methylpimelic acid | 18709 | Not detectable | 17.734 | 179.068 | 174.089 |
| 7 | Phosphatidylethanolamine | Detectable | Not detectable | 15.273 | 750-553 | 767.555 |
| 8. | cytidine diphosphate diacylglycerol (CDP-DG) | Not detectable | Detectable | 13.633 | 824.381 | 841.384 |

On the other side, lipid profile in MCF-7 cells in normal DMSO treated conditions shows the abundance of 17α-(N-acetyl-D-glucosaminyl) estradiol 3-glucosiduronic acid, 2-amino-tetradecanoic acid, 2-methyl-decanoic acid, 3-Methylpimelic acid and Phosphatidylethanolamine over DOX treated cells. In turn, data indicate that MCF-7 cells employ a distinct lipid profile to support growth and proliferation. Another way, DOX treatment forces MCF-7 cells to cell death, possibly by modulating the lipid metabolism that is not linked to growth and proliferation. A study links the lipid profiles of treated leukemic cancer cells with DOX and a distinct lipid profile with an increase in the levels of dodecanamide as predictor of better cell death (39). In recent, mass spectrometry data show the intracellular metabolic adaptations in MCF-7 and drug resistant MCF-7 cells and most of deregulated metabolic pathways include glutathione and amino acid metabolism pathway (40). A metabolomics based investigation elucidate the role of reduced fatty acid metabolism and other metabolic pathways towards apoptotic cell death in MCF-7 cells treated by DOX (41). A low dose of DOX is shown to reprogram metabolic adaptations in MCF-7 cells and may lead to apoptotic cell death (42). In other way. Furthermore, the involvement of deregulated mitochondria and fatty acid metabolism link to the potential role of doxorubicinone to apoptotic cell death in MCF-7 cells in caspase 3 independent cell death pathway.

The presence and biological relevance of CDP-DG is linked with the mitochondria and potentially linked up with apoptosis (35). An interesting study suggests an association between p53, mitochondria and anionic lipid like CDP-DG (19). Within mitochondria, CDP diacylglycerol synthase (CDS) is suggested to catalyze the conversion of phosphatidic acid (PA) to CDP-diacylglycerol from phosphatidic acid that serves as an important component for the synthesis of phosphatidylglycerol, cardiolipin and phosphatidylinositol (PI) (37). In turn, this observation supports our observations that doxorubicinone may disrupt mitochondria and that may lead to the accumulation of CDP-DG and further implicated in the caspase 3 independent apoptotic cell death in MCF-7 cells by DOX. In mitochondrial lipid metabolism, PS is suggested to be synthesized from CDP-DG and serine. There is an enzyme system that involves mitochondrial decarboxylation of PS to generate PE (36, 38, 43). Further, accumulation of substrate such as CDP-choline and other similar structure CDP-DG is suggested to participate in the apoptotic process in cancer cells due to drug treatment (44). In our observation, MCF-7 cells in normal condition show high level of PE. On the other side, DOX treatment to MCF-7 cells leads to decrease in PE level and the accumulation of CDP-DG. Therefore, our finding suggests that DOX induced apoptotic cell death in MCF-7 cells is implicated by the lipid profile alterations in mitochondria and a possible link with the caspase 3 independent cell death pathway.

Sphingolipids that includes sphingosine, sphinganine, and phytosphingosine are known as intracellular metabolites that play a role in growth inhibition and apoptosis in cancer cells (15, 17, 18). Further, evidence indicates that one of sphingolipids i.e. phytosphingosine may participate in the dysfunction of mitochondria and further linked to the induction of apoptosis. However, there is a lack of single direct data that shows the intracellular metabolite identification of phytosphingosine in DOX treated MCF-7 cells and possible link with caspase 3 independent apoptotic cell death in MCF-7 cells.

### Non-lipid intracellular metabolite profiles in DOX treated MCF-7 cells over DMSO control

Besides lipid metabolic reprogramming, metabolic adaptations by cancer cells are seen for polyamine, protein glycosylation and detoxification metabolic pathways (12, 45–50). In this paper, we attempt to highlight the detection and abundance of non-lipid intracellular metabolites in DOX treated MCF-7 cells over DMSO control. Importantly, we claim the use of VTGE assisted purification of intracellular metabolites and further detection by LC-HRMS techniques that facilitated pinpointing key intracellular metabolites that participates in DOX mediated MCF-7 toxicity. An interesting finding convinces that N-acetyl-D-glucosamine, N1-Acetylspermidine and gamma-L-Glutamyl-L-cysteine are potential non-lipid metabolites that are elevated in DOX treated MCF-7 cells compared to DMSO treated cells. Conversely, the intracellular presence of gamma-L-Glutamyl-L-cysteine is higher in the DMSO control MCF-7 cells and almost undetectable in DOX treated MCF-7 cells. This observation may be explained in the sense that due to high load of oxidative stress during DOX mediated cell death, the level of gamma-L-Glutamyl-L-cysteine is exhausted in DOX treated MCF-7 cells. Among various mechanisms that depict the role of DOX in depleting the antioxidant defense system including, glutathione and augmenting the free radical stress in the targeted cancer cells including MCF-7 cells (12).

Furthermore, future investigation is warranted to establish the link between levels of gamma-L-Glutamyl-L-cysteine and DOX mediated sensitivity or resistance towards cancer cells. Among various cellular adaptations in cancer cells, detoxification of reactive oxygen species is considered as a viable option to deal the cellular stress and promote growth. Among potential antioxidant system, glutathione and its precursor γ-Glutamylcysteine are suggested as key intracellular metabolites in cancer cells that support the detoxification of reactive oxygen species (48). Further, earlier findings suggest the elevated expression of gamma-glutamylcysteine synthetase that catalyzes the formation of γ-Glutamylcysteine in case DOX resistant cancer cells (45). With reference to DOX toxicity in MCF-7 cells, we surprisingly notice that in normal condition MCF-7 cells shows the abundance of γ-Glutamylcysteine in the intracellular compartment (Table 2, Figure 8). But due to DOX treatment, MCF-7 cells display an undetectable level of γ-Glutamylcysteine. Interestingly, this finding is again novel in the sense that DOX implicated free radical generation and cellular insults are somehow debilitated in case of DOX treated MCF-7 cells. Therefore, apoptotic cell death in MCF-7 cells by DOX may be partially linked to the reduction in the level of γ-glutamylcysteine and possible inactivation of enzyme γ-glutamylcysteine synthetase that catalyzes the formation of the same. Therefore, VTGE and LC-HRMS based detection of level of γ-Glutamylcysteine in MCF-7 cells provide a new link between DOX and MCF-7 toxicity that is not reported elsewhere with a direct evidence at the level of intracellular metabolite profiling.

**Figure. 8.**
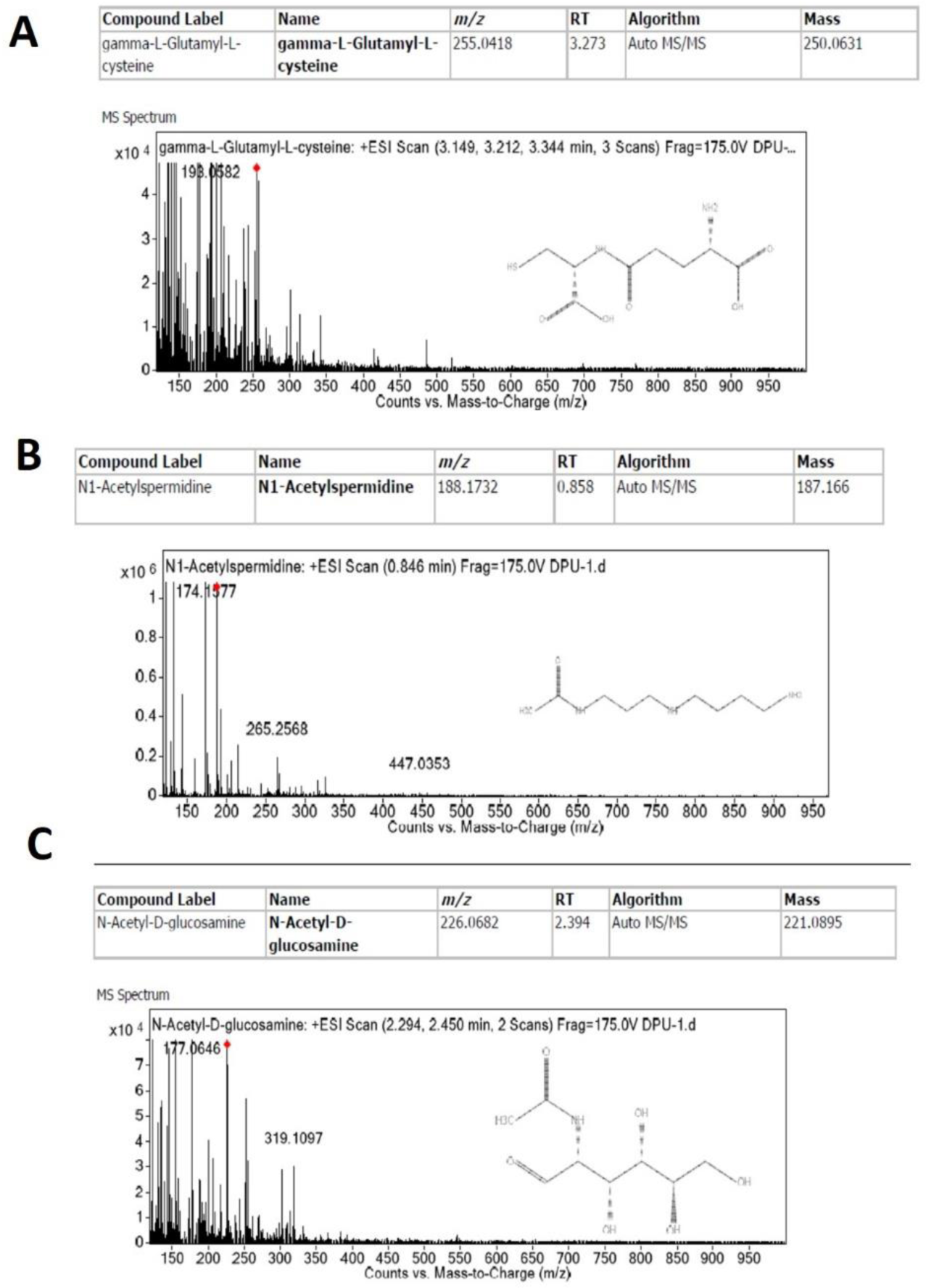
Non-lipid intracellular metabolite alterations between DOX and DMSO treated MCF-7 cells suggest a possible link to DOX mediated toxicity To identify non-lipid metabolite alterations due to DOX, MCF-7 cells were treated as detailed in Figure 2 legend with DOX and DMSO. Further, MCF-7 cells were hypotonically lysed and whole cell lysate were purified by specifically designed VTGE system. Based on LC-HRMS identification and analysis, data suggest that N-acetyl-D-Glucosamine, N1-acetylspermidine and gamma-L-glutamyl-L-cysteine as potential non-lipid metabolite may play a role in DOX treated MCF-7 cells (Figure 8 A-C).

**Table 2:**
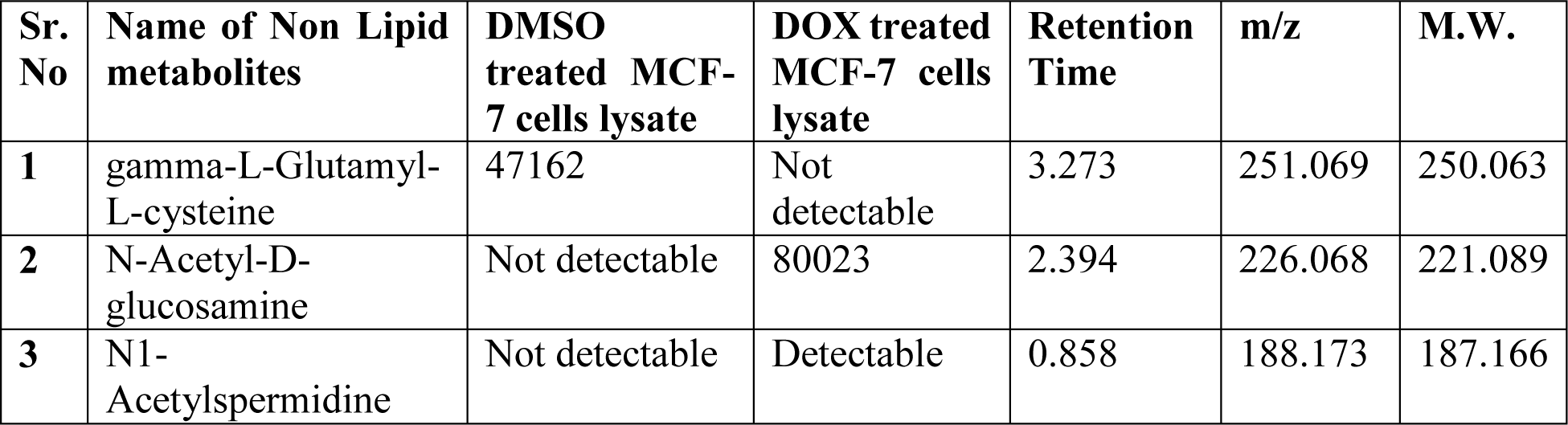
A list of non-lipid intracellular metabolite comparative abundance in DOX over DMSO treated MCF-7 cells.

| Sr. No | Name of Non Lipid metabolites | DMSO treated MCF-7 cells lysate | DOX treated MCF-7 cells lysate | Retention Time | m/z | M.W. |
| --- | --- | --- | --- | --- | --- | --- |
| 1 | gamma-L-Glutamyl-L-cysteine | 47162 | Not detectable | 3.273 | 251.069 | 250.063 |
| 2 | N-Acetyl-D-glucosamine | Not detectable | 80023 | 2.394 | 226.068 | 221.089 |
| 3 | N1-Acetylspermidine | Not detectable | Detectable | 0.858 | 188.173 | 187.166 |

A relatively less known cellular mechanism that includes acetylation of polyamines including spermidine is reported in MCF-7 cells in the context of growth and cellular death (46). Equally, abundance of acetylated polyamines including N (1)-acetylspermidine is suggested to be implicated in the cancer cell growth and death. A very recent paper substantiates partially the changes in the sphingolipid metabolism in MCF-7 cells treated with DOX (20). However, the data are highly limited with reference to DOX mediated intracellular metabolite alterations in the level of acetylated polyamines. Here, the authors propose that abundance of acetylated polyamine is one of the potential mechanisms that may assist in the concerted cellular mechanisms to achieve the cellular death in MCF-7 cells by DOX toxicity. In this paper, our novel observation supports such possibilities where DOX may be able to induce enzymes that catalyze the abundance of acetylation of polyamines as N (1)-acetylspermidine. From the data collected from VTGE and LC-HRMS assisted intracellular metabolite profiling, we notice a high abundance of N(1)-acetylspermidine in DOX trated MCF-7 cells over DMSO control MCF-7 cells (Table 2 and Figure 8). Furthermore, polyamine metabolism is connected to mitochondrial function and cellular energetics that support growth and proliferation (50). Therefore, it appears logical to suggest that DOX treatment in MCF-7 cells may lead to acetylation of polyamines that lead to the arrested growth and toxicity via mitochondrial dysfunction.

Cancer cells are known to engage various enzymes that facilitate glycosylation of key oncogenic proteins using N-acetylglucosamine as a substrate (47, 49). Additionally, functioning of N-acetylglucosamine transferase enzyme is necessary for cellular functions inside cancer cells including MCF-7 cells. In cellular stress and death process, these enzymes are indicated to be dysfunctional and as a consequence, the level of N-acetylglucosamine in the intracellular compartment may be elevated (47, 49). In the same line, we first time report on the high abundance of N-acetylglucosamine in DOX treated MCF-7 cells compared to DMSO control MCF-7 cells. This finding provides a novel information on the possibilities of DOX mediated cell death in MCF-7 cells and glycosylation status.

### Identifications of distinct tripeptide abundance in DOX treated MCF-7 cells

Here, the authors would like to report additional dimension of intracellular metabolite detection as tripeptides in MCF-7 cells treated by DOX and DMSO control. The authors emphasize that tripeptides such as Arg-His-Trp, Cys-Gln-Gln, Glu-Glu-Lys, Pro-Ile-Glu and Gly-Cys-Leu (Figure 9A) are found in the DMSO treated MCF-7 cells (Table 3). Compared to DMSO control, DOX treated MCF-7 cells display distinct identities of tripeptides Arg Trp Pro, Gln Phe Tyr and Lys Glu Glu (Figure 9B) presence in the intracellular compartment. The authors believe that VTGE system and LC-HRMS techniques facilitated the identification of a novel set of tripeptides in DOX and DMSO treated MCF-7 that are not reported in earlier studies centered on DOX and MCF-7 cells. The distinct tripeptide metabolite profiles in DOX treated MCF-7 cells over DMSO control may be explained that the detected tripeptides can be transported by peptide transporter to intracellular compartment to support normal growth conditions. And these tripeptides are originated from serum and therefore, this sounds logical in case of cancer cells that require abnormal growth. On the contrary, the absence of these growth promoting tripeptides in MCF-7 cells treated by DOX may be potential factor in anti-growth effects due to the alterations of membrane properties. Concomitantly, we report on the unique and novel identification of intracellular tripeptides as Arg-Trp-Pro, Gln-Phe-Tyr and Lys-Glu-Glu in MCF-7 cells treated by DOX. At this point, we do not exaggerate on the potential molecular mechanisms behind the evidence on these tripeptides. We also speculate that DOX mediated free radical insults to cellular proteins can lead to the generation of intracellular tripeptides such as Arg-Trp-Pro, Gln-Phe-Tyr and Lys-Glu-Glu. A logical explanation emerges that support the effects of DOX upon MCF-7 to lead intracellular alterations in tripeptides. There are less explored aspects of peptide transporters presence in MCF-7 cells that may participate in pro-growth or drug responsive factors (51, 52). Besides an explanation may support that intracellular peptides in DMSO treated MCF-7 cells can offer protection from the oxidative stress and in turn for growth of MCF-7 cells. On the other hand, absence of oxidative stress protective tripeptides in DOX treated MCF-7 cells may work as one of factors behind responsiveness to DOX mediated cell death. In turn, our findings propose that DOX treated MCF-7 cells showing distinct tripeptide profiles that may support anti-growth and pro-cellular death cellular events. In summary, the authors propose a novel association between intracellular tripeptide metabolites and DOX mediated cell death in MCF-7 cells. In future, such model of molecular mechanisms may be addressed to reveal an additional understanding of DOX mediated cell death in MCF-7 cells.

**Figure. 9.**
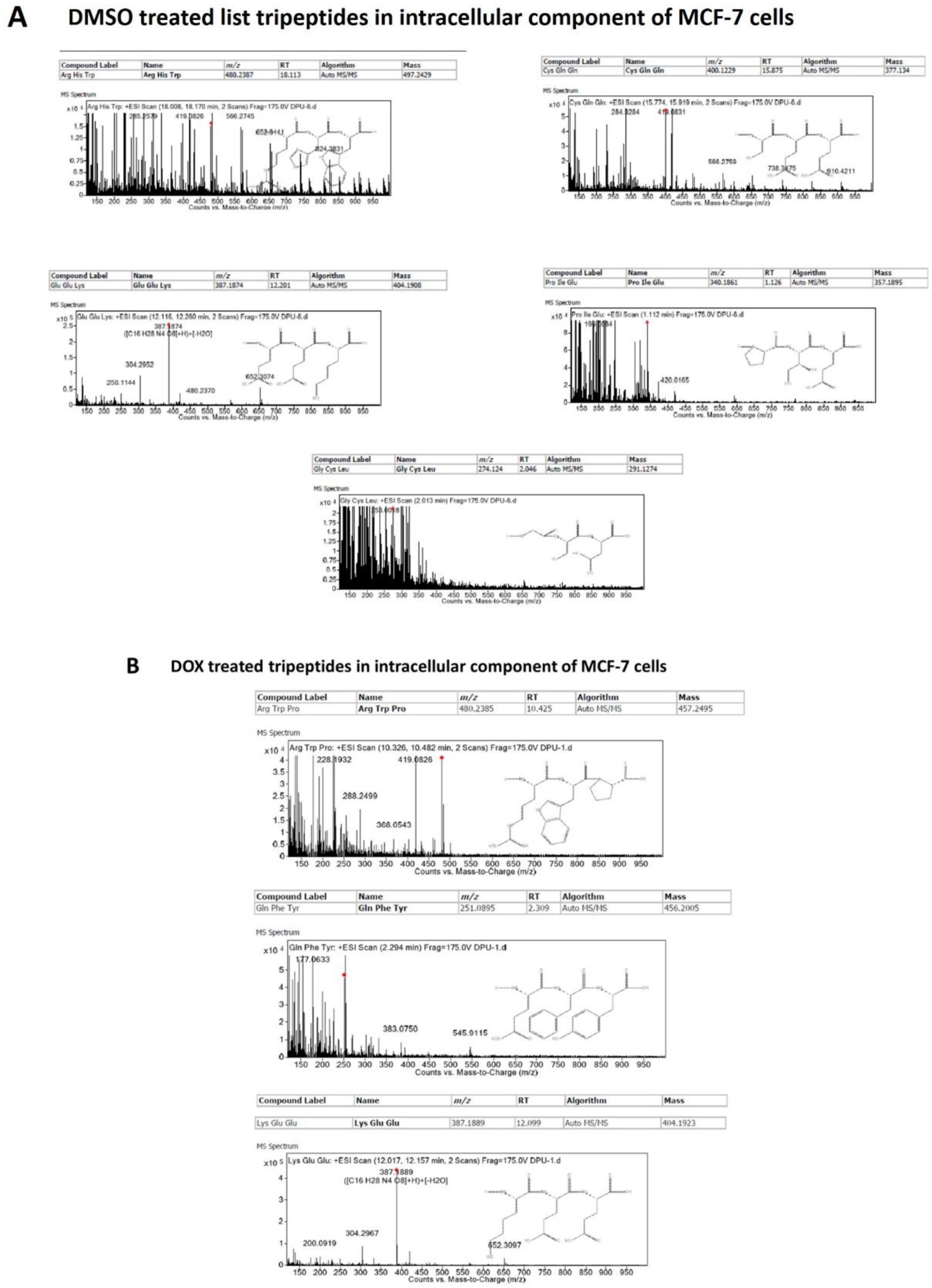
Identification of tripeptide intracellular metabolite profile towards DOX mediated toxicity To identify tripeptide metabolite alterations due to DOX, MCF-7 cells were treated as detailed in Figure 2 legend with DOX and DMSO. Further, MCF-7 cells were hypotonically lysed and whole cell lysate were purified by specifically designed VTGE system. Based on LC-HRMS identification and analysis, data indicate the potential distinctive tripeptide accumulation in the DOX treated MCF-7 cells over DMSO treated MCF-7 cells. Figure 9A depicts on the details the mass ion spectra of DMSO treated MCF-7 cells specific tripeptide metabolites. Figure 9B depicts the mass ion spectra of DOX treated MCF-7 cells specific tripeptide metabolites.

**Table 3:** A list of intracellular tripeptide metabolite in DOX over DMSO treated MCF-7 cells.

| **Sr. No** | **Name of Tripeptides (DMSO treated MCF-7 cells lysate)** | **Retention Time (TIC)** | **Name of Tripeptides (DOX treated MCF-7 cells lysate)** | **Retention Time (TIC)** |
| --- | --- | --- | --- | --- |
| **1.** | Arg His Trp | 18.113 | Arg Trp Pro | 10.425 |
| **2.** | Cys Gln Gln | 15.875 | Gln Phe Tyr | 2.309 |
| **3.** | Glu Glu Lys | 12.201 | Lys Glu Glu | 12.099 |
| **4.** | Pro Ile Glu | 1.126 |  |  |
| **5.** | Gly Cys Leu | 2.046 |  |  |

Taken together, we emphasize on the importance of specifically designed VTGE assisted purification of cellular metabolites and further identification by LC-HRMS technique that helped to identify a novel set of tripeptides in case of DOX treated MCF-7 cells. This data add a new dimension in DOX mediated cell death effects in MCF-7 cells by linking with the abundance of distinct set of intracellular tripeptides. However, further research is warranted to look into this direction.

In summary, this paper reports on a novel and specifically designed method and process to study intracellular metabolic alterations in MCF-7 cells treated with DOX. In this line, identified intracellular metabolites are discussed to support the caspase 3 independent apoptotic cell death with an emphasis on mitochondrial metabolic deregulation and oxidative stress as potential pathways and depicted in Figure 10.

**Figure 10.**
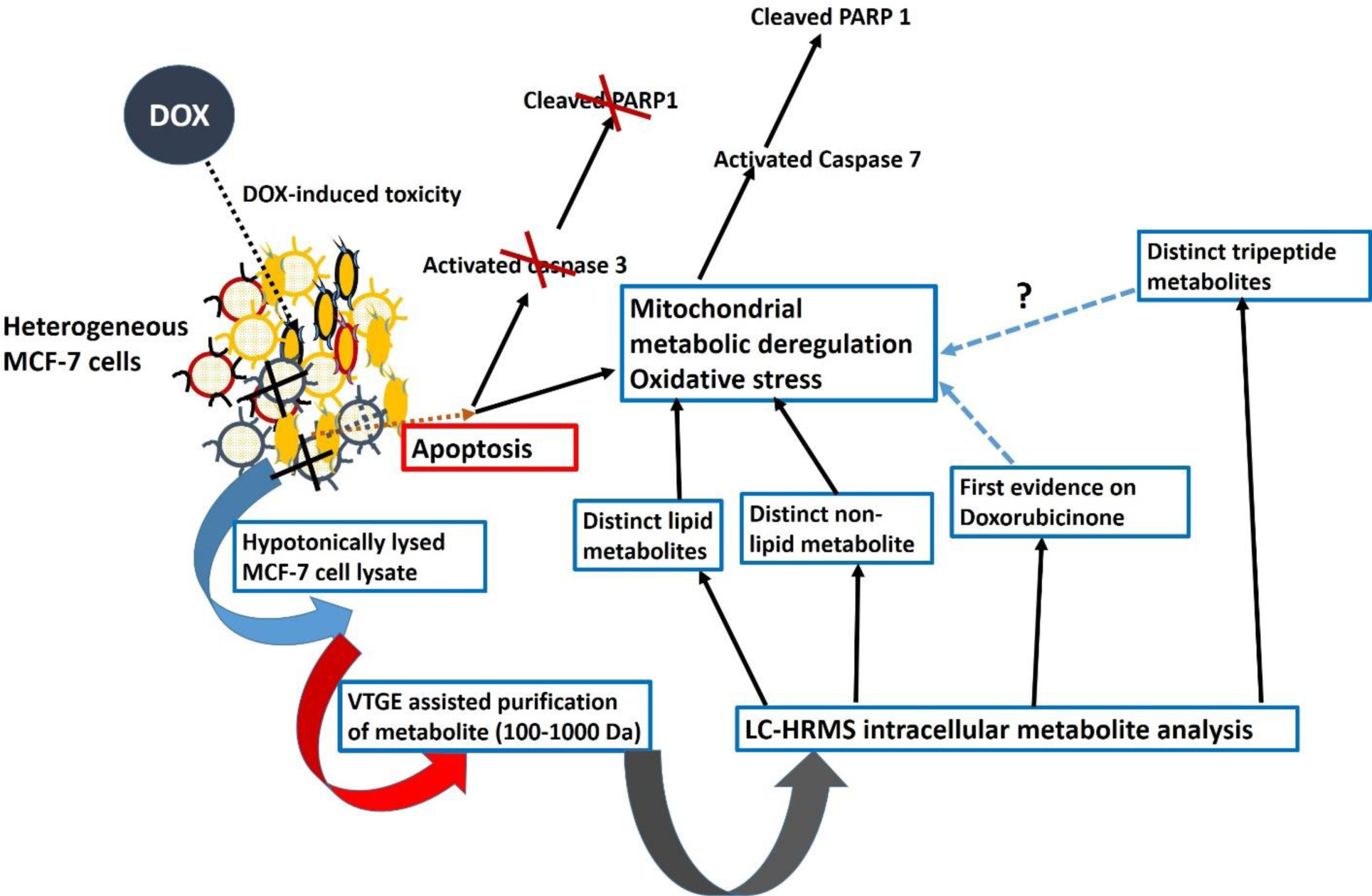
A proposed model of intracellular metabolite adaptations and Dox-induced cell death in MCF-7 cells

## CONCLUSION

The clinical applications of DOX are reported to be useful for both solid and liquid tumors. However, cancer cell plasticity and metabolic adaptation are behind the emergence of DOX mediated drug resistance and the formation of secondary tumors. There are numerous papers that have studied various aspects of DOX and molecular basis of toxicity in cancer cells and normal cells. However, limited findings highlight the intracellular metabolic changes in MCF-7 cells treated by DOX. Interestingly, controversies are quite visible in view of possible cell death pathways either as caspase 3 dependent or caspase 3 independent that may involve mitochondria induced cell death. Our data indicate that DOX is able to bring apoptotic cell death by promoting the cleavage of PARP 1 and caspase 3 independent. Further, not a single paper has shown the intracellular identifications of DOX or its aglycone metabolite in MCF-7 cells treated by DOX. In this direction, we report on a novel methodology and process, including hypotonic preparation of whole cell lysate, followed by VTGE based purification of metabolites and then identification by LC-HRMS. By using these approaches, this paper claims to delineate the precise identification of intracellular metabolite changes in MCF-7 cells due to DOX toxicity. Further, these first time identified lipids (CDP-DG, phytosphingosine, dodecanamide), non-lipids (N-acetyl-D-glucosamine, N1-acetylspermidine and gamma-L-glutamyl-L-cysteine) and tripeptides as intracellular metabolites are discussed to support the potential mechanisms that help to better understand the DOX mediated apoptotic cell death in MCF-7 cells. Besides above findings, the authors conclude that discussed methodologies that analyze intracellular metabolites have potential for compatibility with other cancer cell types and drug regimens to reveal intracellular changes in metabolites towards resistance or susceptibility of a drug.

## ACKNOWLEDGEMENTS

The authors acknowledge financial support from DST-SERB, Government of India, New Delhi, India (SERB/LS-1028/2013) and Dr. D.Y. Patil Vidyapeeth, Pune, India (DPU/05/01/2016).

## CONFLICT OF INTEREST

The authors declare that they have no conflict of interest.

